# Application of slow-controlled release fertilizer coordinates the carbon flow in carbon-nitrogen metabolism to effect rice quality

**DOI:** 10.1101/2023.12.07.570515

**Authors:** Zhengrong Jiang, Qiuli Chen, Dun Liu, Weike Tao, Shen Gao, Jiaqi Li, Chunhao Lin, Meichen Zhu, Yanfeng Ding, Weiwei Li, Ganghua Li, Soulaiman Sakr, Lihong Xue

**Author notes:** **E-mail address:** Zhengrong Jiang, Qiuli Chen, Dun Liu, Weike Tao, Shen Gao, Jiaqi Li, Chunhao Lin, Meichen Zhu, Yanfeng Ding, Weiwei Li, Ganghua Li, Soulaiman Sakr.

## Abstract

Slow-controlled release fertilizers are experiencing a popularity in rice cultivation due to their effectiveness in yield and quality with low environmental costs. However, the underlying mechanism by which these fertilizers regulate grain quality remains inadequately understood. This study investigated the effects of five fertilizer management practices on rice quality in a two-year field experiment: CK, conventional fertilization, and four applications of slow-controlled release fertilizer (UF, urea formaldehyde; SCU, sulfur-coated urea; PCU, polymer-coated urea; BBF, controlled-release bulk blending fertilizer). In 2020 and 2021, the yields of UF and SCU groups showed significant decreases when compared to conventional fertilization, accompanied by a decline in nutritional quality. Additionally, PCU group exhibited poorer cooking and eating qualities. However, BBF group achieved increases in both yield (10.8 t hm^−2^ and 11.0 t hm^−2^) and grain quality reaching the level of CK group. The sufficient nitrogen supply in both the PCU and BBF groups during the grain-filling stage led to a greater capacity for the accumulation of proteins and amino acids in the PCU group compared to starch accumulation. Intriguingly, BBF group showed better carbon-nitrogen metabolism than that of PCU group. The optimal nitrogen supply present in BBF group suitable boosted the synthesis of amino acids involved in the glycolysis/ tricarboxylic acid cycle, thereby effectively coordinating carbon-nitrogen metabolism. The application of the new slow-controlled release fertilizer, BBF, is advantageous in regulating the carbon flow in the carbon-nitrogen metabolism to enhance rice quality.

## 1. Introduction

Rice (*Oryza sativa* L.) is a crucial staple food crop globally, but its production is challenged by two key factors: grain development and grain quality (Thakur et al., 2011; Wang and Peng, 2017; Custodio et al., 2019). With the rapid economic development, there is a growing demand for high-quality rice in developing countries across Asia (Adjao and Staatz, 2015; Concepcion et al., 2015). Grain quality characteristics are essential for determining market value and include factors such as nutritional value, physical appearance, cooking and sensory properties, and milling recovery (Fitzgerald et al., 2009). Fertilization plays a critical role in regulating the grain quality of rice as it affects the content and composition of starch and protein in the grain (Gu et al., 2015). Over the past few decades, conventional fertilizers in rice cultivation face challenges such as low efficiency, high wastage, and labor-intensive application methods, causing environmental pollution and economic losses (Chen et al., 2014). To address these issues, the application of slow-controlled release fertilizers has been proven to be a more efficient method of fertilization, supplying crops with required nutrients in a single basal application throughout the growth period (Naz and Sulaiman, 2016; Qiong et al., 2021). The slow-controlled release fertilizer has a dramatic influence on optimizing grain quality and nitrogen use efficiency, and are widely adopted to minimize economic losses and reduce environmental pollution (Ke et al., 2017; Gao et al., 2021).

A large number of slow-controlled release fertilizers, including urea formaldehyde (UF), sulfur-coated urea (SCU), polymer-coated urea (PCU), and controlled-release bulk blending fertilizer (BBF), have been developed to meet rice’s nutrient demands, enhance yield and quality (Wu et al., 2021). UF and SCU are slow-release fertilizers made from easily decomposable organic materials. UF fertilizer is produced by combining urea with formaldehyde, resulting in quick nitrogen release during the early growth stage but limited release during the middle and late stages (Shaviv, 2001). In contrast, SCU fertilizer involves coating urea granules with a layer of sulfur, enabling the nutrients to be slowly released over time; however, the release rate is insufficient during the middle and late stages (Xu et al., 2016). PCU and BBF are two types of controlled-release fertilizers that provide a regulated-release of nutrients to plant. The PCU fertilizer is an enhanced version of SCU, designed to minimize the impact of microorganisms on nutrient release through coating urea granules with a polymer material. BBF is created by blending various types of slow-release and controlled-release fertilizers, allowing for a consistent release of nitrogen nutrients throughout the entire rice growth cycle (Ye et al., 2013). Given the growing demand for grain quality and efficient fertilizer use, further research is needed to examine the relationship between grain quality and the application of slow-controlled release fertilizers.

Grain filling is vital for rice quality and is highly influenced by fertilizer use. Accumulation of lipids, starch, and storage proteins during grain filling stage is crucial for grain quality, and is strongly influenced by nitrogen availability (Jiang et al., 2021; Wang et al., 2021b; Xu et al., 2021). The improper use of fertilizer can negatively impact the physical appearance of rice grain, leading to the formation of chalkiness as a result of an unbalanced carbon-nitrogen metabolism (Zhao et al., 2019; Wang et al., 2021b). The carbon-to-nitrogen (C: N) balance in rice grains refers to the optimal ratio of protein and starch in the grain, which is critical for determining the nutritional value and other rice qualities (Dou et al., 2017). The changes in nitrogen and carbon supply can alter numerous central metabolites involved in carbon and nitrogen metabolism in parallel (Gao et al., 2020). To support the synthesis of amino acids and ensure adequate growth in rice grain, an appropriate nitrogen content is needed to drive the production of carbon skeletons and provide enough amino acids (Gutiérrez et al., 2007; Wang et al., 2022a).

The inorganic nitrogen of soil is absorbed by roots and can then be assimilated into glutamine and glutamate in grain, which serves as a source of organic nitrogen (Huang et al., 2021). Glutamine synthetase (GS), encoded by *OsGS2*, is a crucial enzyme for assimilating NH_4_ ^+^ in rice. Maintaining a stable GS level is essential for maintaining the carbon-nitrogen balance during rice growth (Bao et al., 2015). The key enzymes of the GS and GOGAT (glutamate synthase) cycle play key roles in nitrogen assimilation of rice grain (Luo et al., 2018; Poucet et al., 2021). Additionally, nitrogen assimilation requires energy and carbon skeletons, which are derived from sucrose, glucose, other glycolysis-derived carbohydrates (pyruvate) and the tricarboxylic acid (TCA)-derived organic acids (eg., cetoglutarate and oxaloacetate) (Huang et al., 2021). The carbon and nitrogen metabolites are also involved in the oxidative pentose phosphate pathway (OPPP), which is crucial for generating the energy and reducing power required for rice grain growth (Li et al., 2016; Yang et al., 2016; Wang et al., 2021a). The TCA cycle is essential for providing 2-oxoglutarate, which is necessary for the production of glutamate and glutamine involved in nitrogen assimilation (Nunes-Nesi et al., 2010). The intermediates of the glycolysis pathway, like pyruvate, play an important role in the TCA cycle and nitrogen assimilation in rice (Masumoto et al., 2010).

The primary product of photosynthesis, sucrose, plays a crucial role in supplying the carbon skeletons required for synthesizing a variety of macromolecules, such as proteins, nucleic acids, lipids, and starch (Jiang et al., 2021). This is facilitated by the transport pathways of *OsCIN4* (invertase) and *OsSUT2* (sucrose transporter) during grain filling (Wang et al., 2008; Deng et al., 2021). The major function of SuSase is to breakdown sucrose to supply carbohydrates for the process of OPPP and glycolysis/TCA cycle, and the enzymes AGPase, GBSS and SBE play major roles in catalyzing the synthesis of starch to promote grain filling (Wang et al., 2021a; Xu et al., 2022). However, the mechanisms linking carbon and nitrogen metabolism to rice grain growth under the influence of slow-controlled release fertilizer are not yet fully understood.

In this study, a two-year field experiment was conducted to examine the impact of various slow-controlled release fertilizers on grain quality of rice. To gain a deeper understanding of the nitrogen assimilation process in rice grain under the influence of slow-controlled release fertilizers, we also analyzed the underlying carbon-nitrogen mechanisms during grain filling. Our findings will provide a solid foundation for improving the quality and yield of rice through the use of slow-controlled release fertilizers.

## 2. Materials and methods

### 2.1 Study site and N sources

Field experiments were conducted over two rice growing seasons (2020 and 2021) in Yanling Town of Danyang city, Jiangsu Province, China (31°54′31″N, 119′28’21″E). The meteorological data came from the meteorological station (Watch Dog 2900ET, SPECTRUM, USA) installed 100 m from the experimental station (Figure.S1). The soil properties were classified as Orthic Acrisol (Falsone, 2013). The soil had the following characteristics: pH = 6.4, organic matter = 18.32 g·kg^−1^, total N = 1.27 g·kg^−1^, Olsens-P = 16.85 mg·kg^−1^ and NH_4_OAc-extractable K = 139.63 mg·kg^−1^. The slow-controlled release fertilizers used in this study were: UF (35% N), SCU (37% N), PCU (43% N), and BBF (40% N). The UF, SCU, PCU and BBF were provided by Hanfeng Slow Release Fertilizer Co., Ltd. (Jiangsu, China). Conventional urea (46% N) fertilizers were used as CK group for comparison.

### 2.2 Experimental design

For analyzing effects of different controlled-release fertilizers on grain quality of rice, the experiment was conducted by rice cultivation of Ningjing 8^th^ (*Oryza sativa* L.), a high-yielding japonica rice cultivar bred by Nanjing Agricultural University (NAU). This experiment included treatments as follows: a conventional fertilization (CK, four spilt applications of urea), and a single basal application of slow-controlled release fertilizer (SCU, PCU, UF, and BBF). The N release rate of the slow-controlled release fertilizer was shown in Figure. 1, which was determined by buried bag method (Yang et al., 2011). The conventional urea fertilization was split among basic fertilizer, tiller fertilizer, spikelet-promoting fertilizer, and spikelet-developing fertilizer. Each treatment was applied nitrogen at 210 kg·ha^−1^, and for all treatments, P and K fertilizers were applied as basal dressings at 135 kg·ha^−1^ (P_2_O_5_) and 216 kg·ha^−1^ (K_2_O). The field plots, measuring 7.2 m × 20 m, were arranged in a randomized complete block design with three replicates per treatment. In each replicate, a total of approximately 1600 panicles with similar growth patterns were labeled and monitored for their flowering date. This was done in September, when about 50% panicles had fully emerged from the flag leaf sheath.

**Figure 1.**
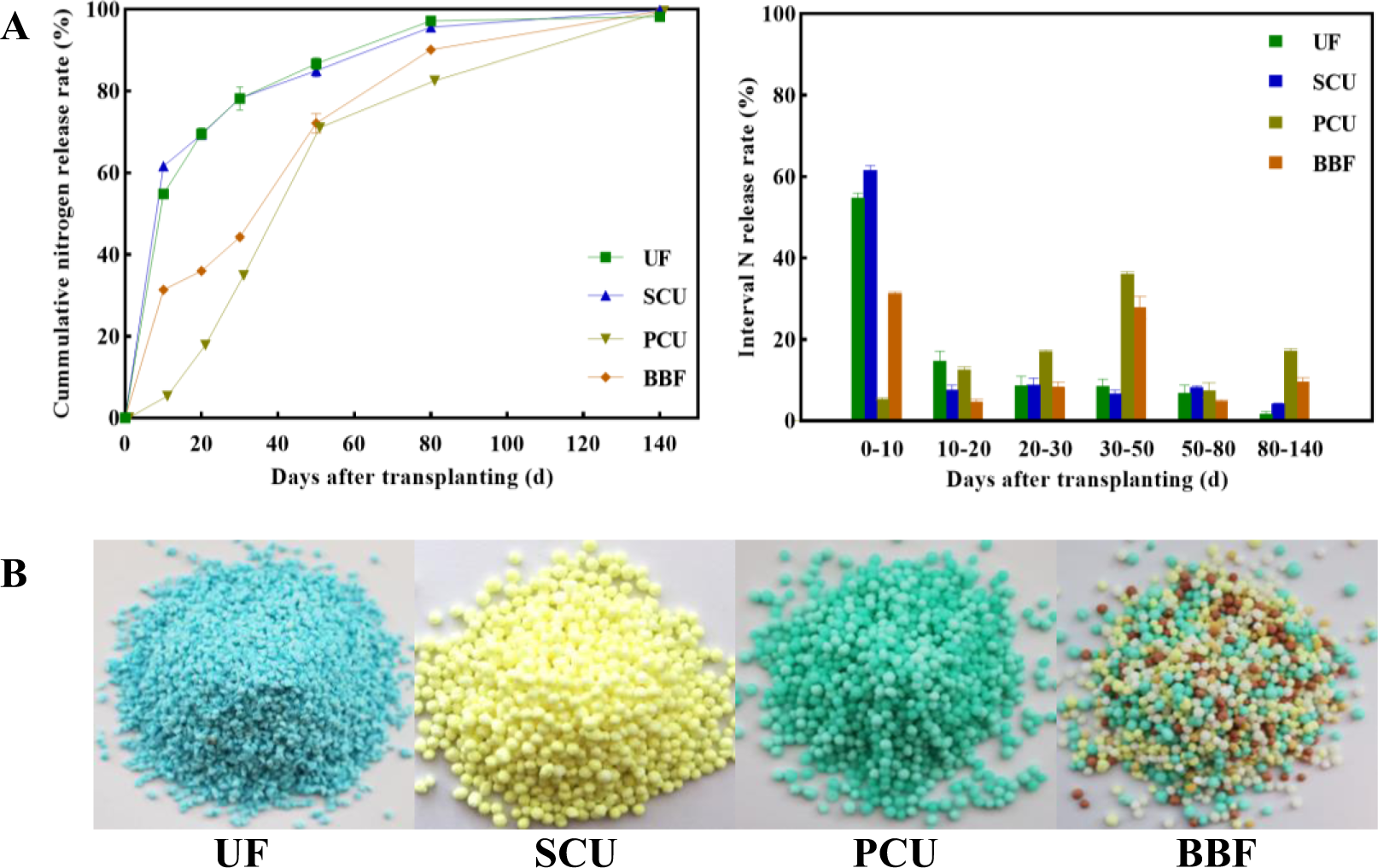
Nitrogen release rate (A) and appearance feature (B) of different types of slow-controlled release fertilizer. CK, conventional fertilization with four spilt applications of urea; UF, urea formaldehyde; SCU, sulfur-coated urea; PCU, polymer-coated urea; BBF, controlled-release bulk blending fertilizer.

### 2.3 Sampling and analysis

#### 2.3.1 Harvesting and grain quality measurements

At maturity stage, all necessary plants (excluding edge-row plants) from each plot were harvested. The grain yield was determined by multiplying each yield component, which had been measured after removing impurities and adjusting moisture content to 14.5%. Before harvesting, the number of effective tillers per hill was determined by selecting 50 plants from each plot. At maturity, approximately 30 marked plants were used to analyze the 1000-grain weight, seed setting rate, and grain number per panicle.

At maturity, the rice grain quality was analyzed. For rice milling quality, the brown rice yield and milled yield were analyzed according to the method of rice measurement standards (NY147-88; Ministry of Agriculture, PR China, 1988). To test the appearance quality of rice, the chalk characteristics of brown rice were determined by the cleanliness test-bed according to Tang et al. (2019). Rice cooking and eating quality was analyzed by the Rapid Visco Analyser (RVA, Starchmaster 2, Perten Instruments of Australia Pty Ltd., Sydney, New South Wales, Australia) to obtain profile characteristics according to Standard Method AACC61-02. Viscous profile characteristics of RVA were expressed as peak viscosity, hot viscosity, cool viscosity, breakdown viscosity, setback (difference between final viscosity and peak viscosity), and consistency. Rice taste characteristics were analyzed by an STA-1A device (Satake Corp., Hiroshima, Japan) according to the method of Jin et al. (2004).

#### 2.3.2 Grain weight and grain growth rate

To analyze the grain filling process, 90 tagged panicles of each replicate plot were sampled every 7 days post anthesis (DPA) from 7 to 42 DPA (7, 14, 21, 28, 35, and 42 DPA). All spikelets were deactivated at 105°C for 60 min and then dried at 80°C to determine the grain dry weight (DW). Grain filling processes were fit to Richards’s growth equation (Richards, 1959).

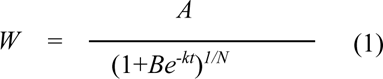

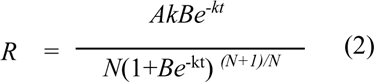

Grain weight (mg), W; Grain filling rate, R; Final grain weight (mg), A; Time after anthesis (days), t; and B, k, and N are coefficients established from the regression of the equation.

#### 2.3.3 Analyze of sugar, starch and storage protein composition

To analyze the sugar and starch content, all samples were first ground into a fine powder and then passed through a 100-mesh sieve. The sugar and starch were extracted using a modified version of the method described by Yang et al. (2001). Approximately 100 mg of the ground sample was mixed with 8 mL of 80% (v/v) ethanol at 80°C for 30 min. After cooling, the tube was centrifuged at 5000 g for 15 min, and the supernatant was collected. The extraction process was repeated three times, and the sugar extract was diluted with distilled water to a final volume of 50 mL. After the sugar extraction, the residues in the tubes were dried at 80°C to extract starch using HCLO_4_ according to Yang’s method. The amylose and amylopectin contents were measured using the Megazyme Amylose/Amylopectin Assay Kit (Megazyme International Ireland Ltd, Bray, Ireland).

The total protein content was measured using an AutoKjeldahl Unit K-370 nitrogen analyzer (Büchi Labortechnik, Flawil, Switzerland) with AACC International Approved Method 46-30.01. Different protein components including albumin, globulin, prolamin, and glutelin were extracted in a stepwise manner using different solvents such as distilled water, dilute hydrochloric acid, ethanol, and dilute alkali. The glutelin content was determined using the biuret colorimetric method, while the remaining protein components were measured using the Coomassie brilliant blue colorimetric method.

#### 2.3.4 Measurements of amino acid composition

To hydrolyze the rice powder samples, approximately 10 mg of the sample was mixed with 1 mL of 6 N HCl (Sigma, USA) in a 2 mL screw-cap tube. After heating at 110°C for 24 h, the samples were further treated for 6 h at 65°C. The resulting residue was dissolved in 1 mL of Na-S™ buffer, and the mixture was centrifuged at 1600 × g for 10 min. The supernatant was filtered through a 0.45 μm nylon membrane syringe filter (Pall Life Sciences, USA), and about 10 nmol of L-(+)-norleucine (Wako Pure Chemicals, Japan) was added. Amino acid analysis was performed using a Hitachi L-8900 amino acid analyzer (Hitachi Corp, Japan) based on the national standard of the People’s Republic of China (Ministry of Agriculture PR China, 1988). The amino acids tested included aspartic acid (Asp), glutamic acid + glutamine (Glu), glycine (Gly), histidine (His), isoleucine (Ile), leucine (Leu), lysine (Lys), phenylalanine (Phe), serine (Ser), threonine (Thr), tyrosine (Tyr), and valine (Val). Total amino acid content was the sum of all amino acid content in the same period. All experiments were conducted at least three biological replicates in each sample, and the HPLC data were normalized to the level of L-(+)-norleucine per sample.

#### 2.3.5 Enzyme extraction and analysis

For enzyme extraction, the grains collected at 7, 14, 21, 28, 35, and 42 DPA were ground into fine powder in liquid nitrogen. The activities of several enzymes including sucrose synthase (SuSase), adenine diphosphoglucose pyrophosphorylase (AGPase), starch branching enzyme (SBE), and granule-bound starch synthase (GBSS) were determined using the method described by Yang et al. (2003). In addition, the activities of glutamine synthetase (GS) and glutamate synthetase (GOGAT) were analyzed according to the protocols outlined by Tang et al. (2018).

#### 2.3.6 RNA extraction and qRT-PCR

Total RNA was extracted from frozen rice grains using a previously described method (Yasuda et al., 2005). Gene transcription levels of the relevant genes were analyzed using RNase-free DNase I treatment, cDNA synthesis, and quantitative real-time polymerase chain reaction (qRT-PCR). The RNA-prep pure PLANT Kit (DP432, Tiangen Biotek, Beijing, China) was used to isolate total RNA from the rice grains, which was then reverse-transcribed into first-strand cDNA using the Prime-Script-TM RT Reagent Kit (RR036, Takara, Kyoto, Japan). The quantitative real-time polymerase chain reaction was performed using the ABI 7300 sequencer and SYBR Premix Ex Taq-TM (RR420, Takara, Kyoto, Japan) according to the manufacturer’s protocol. Each sample was replicated three times to test the expression of genes (*OsSUT2*, *OsCIN4*, and *OsGS2*). The primers used in this study are listed in the Supporting Information (Table S1).

### 2.4 Statistical analysis

Excel 2019 (Microsoft Corp. Redmond, WA, USA), SPSS version 20.0. (SPSS Statistics, SPSS Inc., Chicago, USA), and Origin 2021 (OriginLab, Northampton, MA, USA) were used for data visualization. Data was performed the analysis by using two-way analysis of variance (ANOVA) and principal component analysis (PCA). Means were compared based on the least significant difference at *P* = 0.05 (LSD_0.05_).

## 3. Results

### 3.1 Influence of slow-controlled release fertilizer on rice yield

Compared to the conventional fertilizer’s yield (10.2 t hm^−2^ and 11.2 t hm^−2^) in 2020 and 2021, the rice yields decreased to 8.4 t hm^−2^ and 9.7 t hm^−2^ after application of urea formaldehyde fertilizer (UF), in consistent with the decline in yield (9.6 t hm^−2^ and 8.6 t hm^−2^) of sulfur-coated urea fertilizer (SCU) (Table 1). The groups of PCU and BBF showed similar or even higher yield parameters compared with CK group (Table 1). In 2020, the yield of the BBF group reached 10.8 t hm^−2^, marking the highest yield among all types of fertilizers. Similarly, in 2021, the BBF yield increased to 11.0 t hm^−2^, significantly surpassing the yields of the UF and SCU groups.

**Table 1.** Agronomic traits at the maturity stage under different types of slow-controlled release fertilizer in 2020 and 2021.

| Year | Fertilizer type | Yield (t hm<sup>-2</sup>) | Panicles (No./m<sup>2</sup>) | Spikelets (No./panicle) | Spikelets (No./hm<sup>2</sup>) | Seed setting rate (%) | Grain weight (mg) |
| --- | --- | --- | --- | --- | --- | --- | --- |
| 2020 | CK | 10.2ab | 314.4ab | 129.0a | 4.1a | 93.9a | 26.9c |
|  | UF | 8.4c | 286.2c | 112.3b | 3.2c | 94.9a | 27.6ab |
|  | SCU | 9.6b | 305.5abc | 117.5b | 3.6bc | 95.4a | 28.0a |
|  | PCU | 10.2ab | 293.3bc | 131.8a | 3.9ab | 94.9a | 27.8a |
|  | BBF | 10.8a | 325.3a | 131.2a | 4.3a | 93.6a | 27.2bc |
| 2021 | CK | 11.2a | 323.0a | 137.9a | 4.4a | 96.8a | 26.1c |
|  | UF | 9.7b | 335.7a | 110.5c | 3.7b | 93.6a | 28.3a |
|  | SCU | 8.6b | 343.3a | 94.2c | 3.2c | 94.9a | 28.1ab |
|  | PCU | 11.1a | 368.3a | 116.0abc | 4.3a | 96.3a | 27.2b |
|  | BBF | 11.0a | 355.7a | 117.3ab | 4.2a | 95.6a | 27.5ab |
| F-Value | T | ** | ns | ** | ** | ns | ** |
|  | Y | ** | ** | * | * | ns | ns |
|  | T×Y | ** | ns | ns | ** | ns | * |

CK, conventional fertilization with four spilt applications of urea; UF, urea formaldehyde; SCU, sulfur-coated urea; PCU, polymer-coated urea; BBF, controlled-release bulk blending fertilizer; Values with different letters in the same column are significantly different at $P \leq 0.05$ by the LSD test; \*\*and\*significant at 0.01and 0.05 probability level respectively; ns, not significant.

The application of UF, SCU, PCU, and BBF all resulted in a significant improvement in grain weight compared with the CK group in 2019 and 2020 (Table 1). The differences in grain weight among the various fertilizer group were even more conspicuous in 2021, particularly with the application of BBF and UF significantly promoting grain weight accumulation in comparison to CK group (Figure 2). Concurrently, the grain filling rate of BBF and UF groups was significantly higher than that of other treatments at the middle grain-filling stage (14 DPA). These results demonstrate that the use of suitable slow-controlled release fertilizers, such as BBF, can enhance grain growth and rice yield.

**Figure 2.**
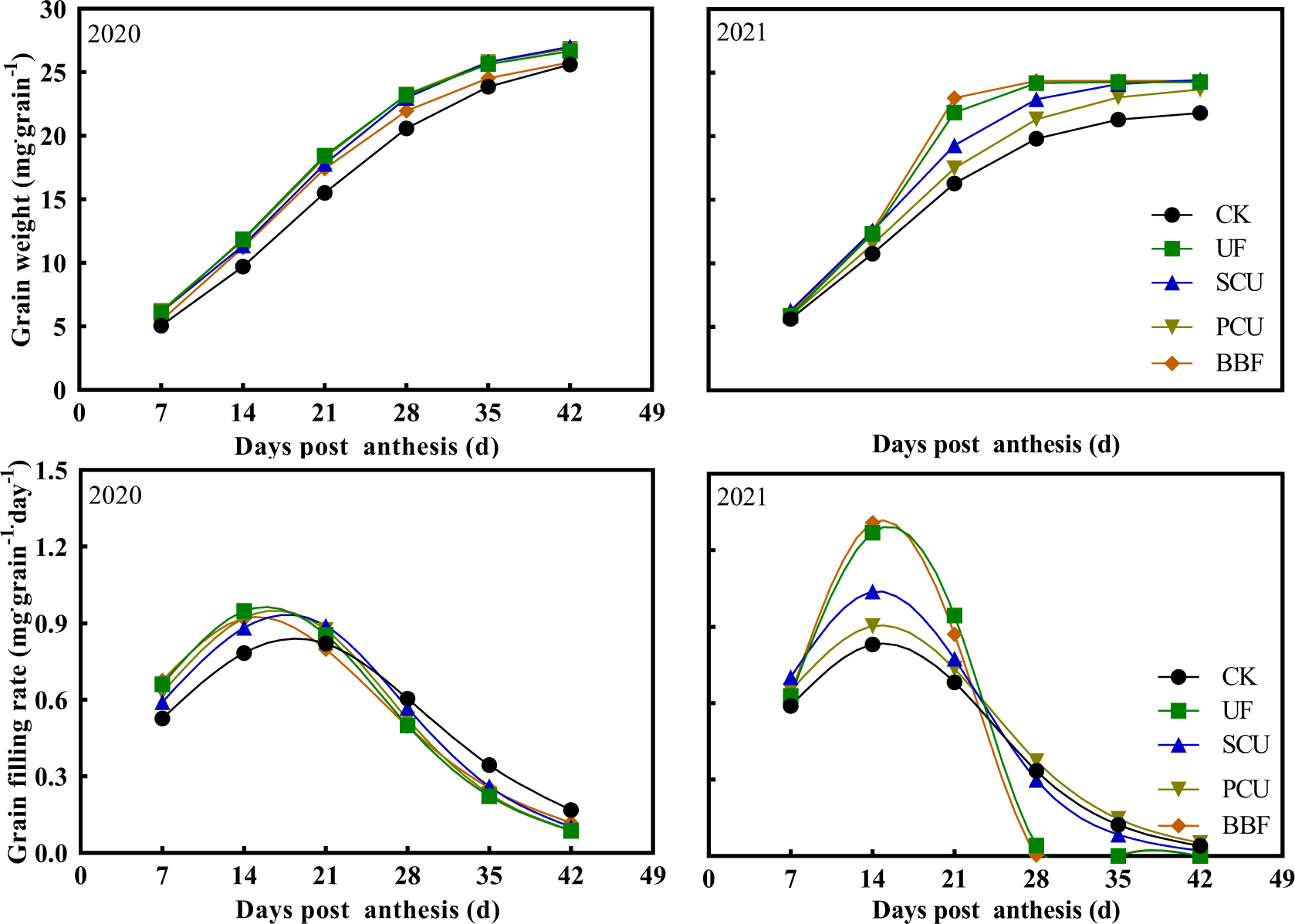
Dynamics of rice grain growth after the heading stage under different types of slow-controlled release fertilizer in 2020 and 2021. CK, conventional fertilization with four spilt applications of urea; UF, urea formaldehyde; SCU, sulfur-coated urea; PCU, polymer-coated urea; BBF, controlled-release bulk blending fertilizer.

### 3.2 Effects of slow-controlled release fertilizer on rice quality

In 2020 and 2021, the application of BBF significantly improved the physical appearance, cooking and eating qualities, as well as the nutritional value of rice, reaching levels comparable to those of the CK group (Tables 2-3). Nevertheless, the taste values for the PCU group were noticeably lower at 65 and 73 in 2020 and 2021, respectively, compared to the CK group’s values of 72.7 and 75 (Table 2). Additionally, the PCU group displayed notably lower cool viscosity and breakdown than the CK group (Table 2). The taste value is critical for cooking and eating qualities and is tightly correlated with the RVA value (Kesarwani et al., 2016). These results indicate that the poor taste value and RVA value resulted in poor cooking and eating qualities in PCU group. The composition of protein and amino acids greatly contributes to the nutritional value of rice grain (Amagliani et al., 2017). Compared to the CK group, the application of UF and SCU resulted in a lower content of nitrogen metabolites in rice grains (Table 3), indicating that the use of UF and SCU was unfavorable to the nutritional value of rice grain.

**Table 2.** Effects of different types of slow-controlled release fertilizer on grain quality of rice in 2020 and 2021.

| Year | Fertilizer type | Brown rice rate (%) | Milled rice rate (%) | Head rice rate (%) | Chalky kernel rate (%) | Chalky area (%) | Chalkiness (%) | Taste value |  |
| --- | --- | --- | --- | --- | --- | --- | --- | --- | --- |
| 2020 | CK | 85.06a | 74.91a | 70.19a | 39.67a | 22.52ab | 8.88ab | 72.7a |  |
|  | UF | 84.23b | 74.5a | 69.22a | 34.17a | 20.80b | 7.17b | 73.0a |  |
|  | SCU | 84.63ab | 74.78a | 69.8a | 34.83a | 21.72ab | 7.61ab | 72.3a |  |
|  | PCU | 84.9a | 75.01a | 70.45a | 41.00a | 26.00a | 10.64a | 65.0b |  |
|  | BBF | 84.8a | 75.3a | 70.46a | 34.83a | 20.46b | 7.11b | 71.3a |  |
| 2021 | CK | 84.10a | 72.53a | 68.47a | 37.46ab | 24.65b | 9.18b | 75.00a |  |
|  | UF | 83.82a | 73.46a | 69.25a | 32.99b | 21.71c | 7.15d | 76.67a |  |
|  | SCU | 83.72a | 74.23a | 71.64a | 36.98ab | 22.56c | 8.34bc | 76.00a |  |
|  | PCU | 83.46a | 72.79a | 69.31a | 42.41a | 27.07a | 11.47a | 73.00b |  |
|  | BBF | 83.8a | 73.64a | 69.97a | 34.80a | 22.31c | 7.75cd | 76.33a |  |
| Year | Fertilizer type | Peak viscosity (cP) | Hot viscosity (cP) | Cool viscosity (cP) | Breakdown (cP) | Setback (cP) | Consistency (cP) | Pasting temperature (°C) | Peak time (min) |
| 2020 | CK | 3270b | 2249ab | 2820ab | 1021b | 450bc | 570ab | 72.1a | 6.6a |
|  | UF | 3390a | 2187b | 2858ab | 1204a | 532a | 671a | 71.9a | 6.4a |
|  | SCU | 3342ab | 2209ab | 2913a | 1133ab | 429c | 704a | 71.9a | 6.5a |
|  | PCU | 3170c | 2298a | 2745c | 872c | 425c | 447b | 72.1a | 6.6a |
|  | BBF | 3277b | 2194b | 2773bc | 1082b | 504ab | 579ab | 72.0a | 6.4a |
| 2021 | CK | 3527c | 2372a | 3051a | 1355d | 476d | 679a | 75.0a | 6.2a |
|  | UF | 3695a | 2334a | 3060a | 1559a | 601c | 726a | 74.2a | 6.0a |
|  | SCU | 3711a | 2281a | 2991a | 1464b | 720a | 711a | 73.9a | 6.1a |
|  | PCU | 3529c | 2152b | 2837b | 1303e | 692ab | 685a | 74.4a | 6.1a |
|  | BBF | 3628b | 2302a | 3007a | 1403c | 622bc | 705a | 74.2a | 6.1a |

CK, conventional fertilization with four spilt applications of urea; UF, urea formaldehyde; SCU, sulfur-coated urea; PCU, polymer-coated urea; BBF, controlled-release bulk blending fertilizer; Values with different letters in the same column are significantly different with $P \leq 0.05$ .

**Table 3.**
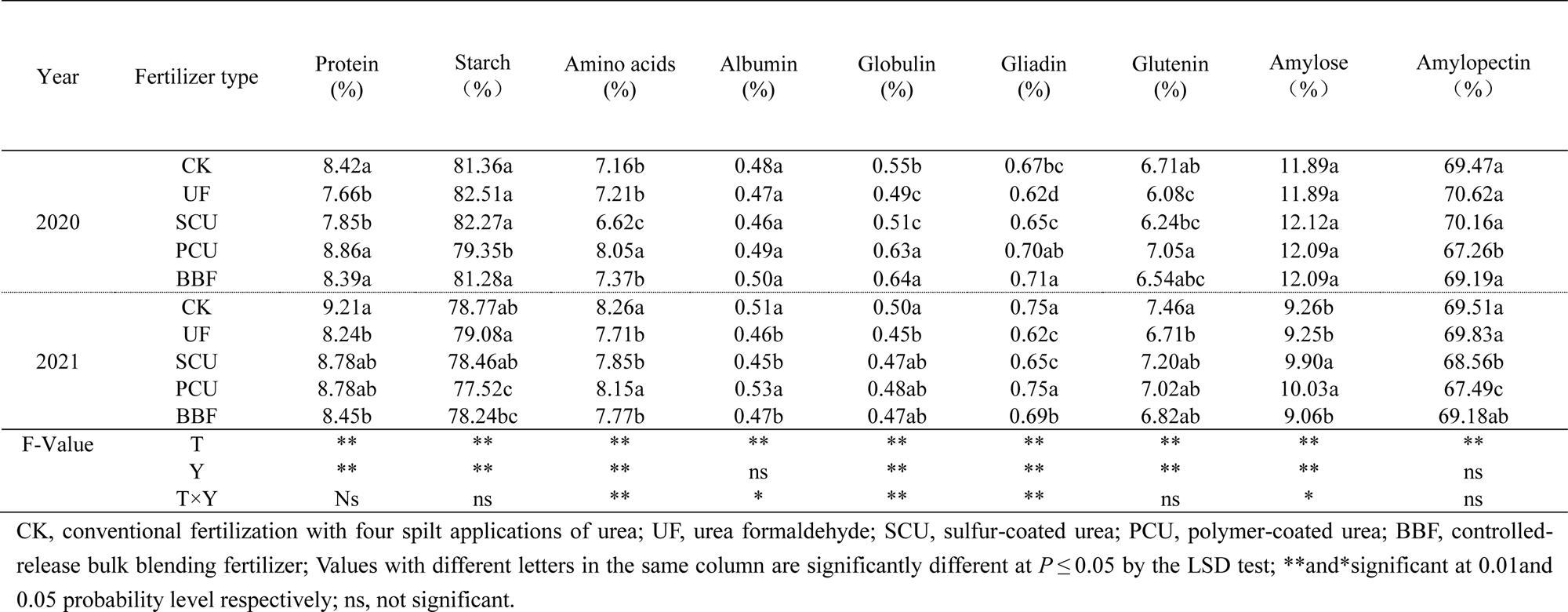
Carbon and nitrogen metabolites at the maturity stage under different types of slow-controlled release fertilization in 2020 and 2021.

### 3.3 Carbon metabolism of rice grain

The carbon metabolism pathway plays a crucial role in rice quality by facilitating the synthesis of starch and protein (Dai et al., 2013; Jiang et al., 2021). The sugar content of all slow-controlled release fertilizer treatments reached or even exceeded the level of CK group at grain-filling stage in 2021 (Figure. 3), while the PCU group showed significantly lower accumulation of starch and amylopectin compared to the CK treatment during grain-filling period (Table 3, Figure. 3). The application of slow-controlled release fertilizers obviously affected the activity of key enzymes involved in carbon-nitrogen metabolism in rice grains (Table S2). The PCU group did not exhibit the lowest enzyme activities related to carbon metabolism (SuSase, AGPase, GBSS, and SBE) in grains among all treatments, but the UF group demonstrated lower enzyme activities during grain filling (Figure. 4). However, the starch content in the UF group was not the lowest (Figure. 4). The carbon metabolite content and the related key enzyme activity of BBF were neither the highest nor the lowest among these treatments (Figures. 3-4).

**Figure 3.**
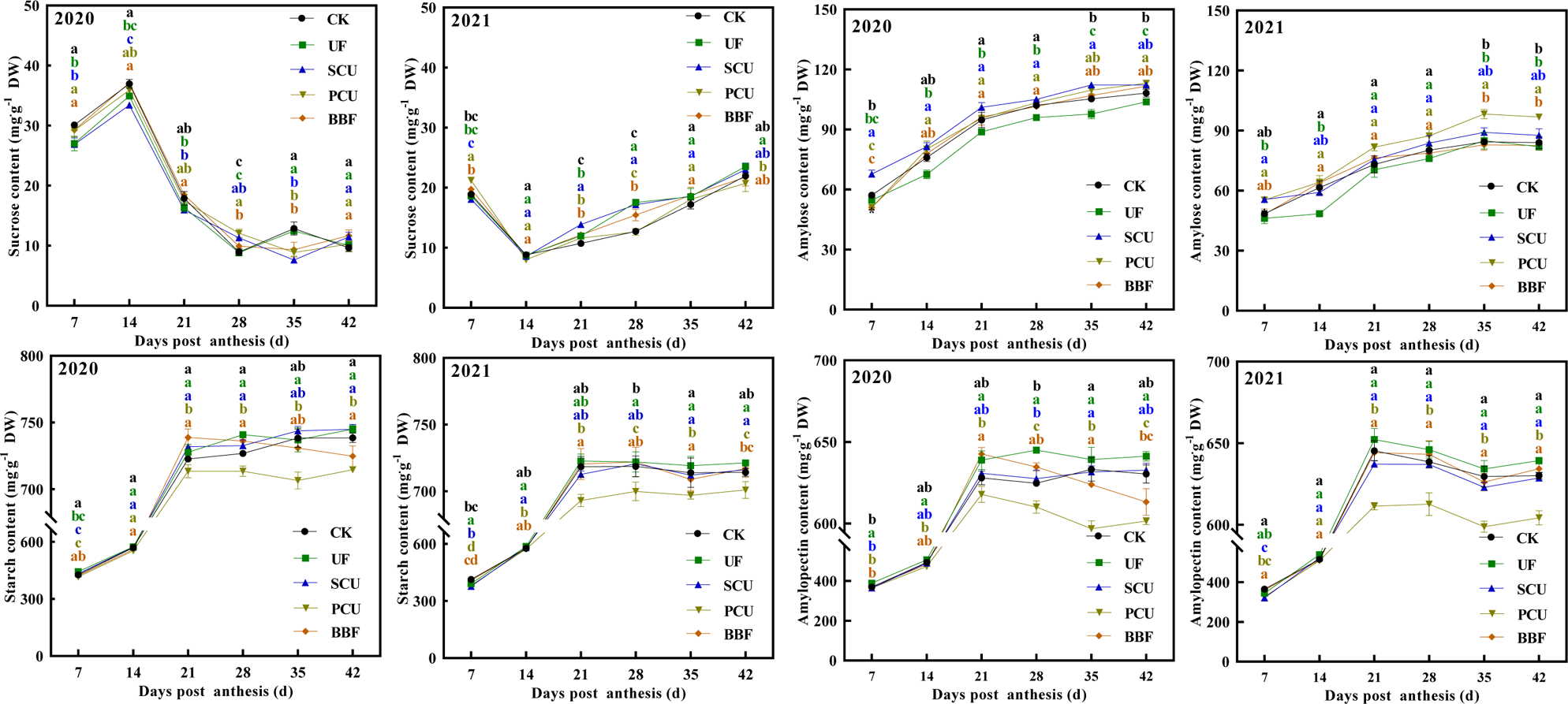
Effects of different types of slow-controlled release fertilizer on the changes in carbohydrates in rice grain during the filling stage in 2020 and 2021. CK, conventional fertilization with four spilt applications of urea; UF, urea formaldehyde; SCU, sulfur-coated urea; PCU, polymer-coated urea; BBF, controlled-release bulk blending fertilizer; Significant differences at each time point are indicated by different letters (*P* < 0.05) as determined by Duncan’s test.

**Figure 4.**
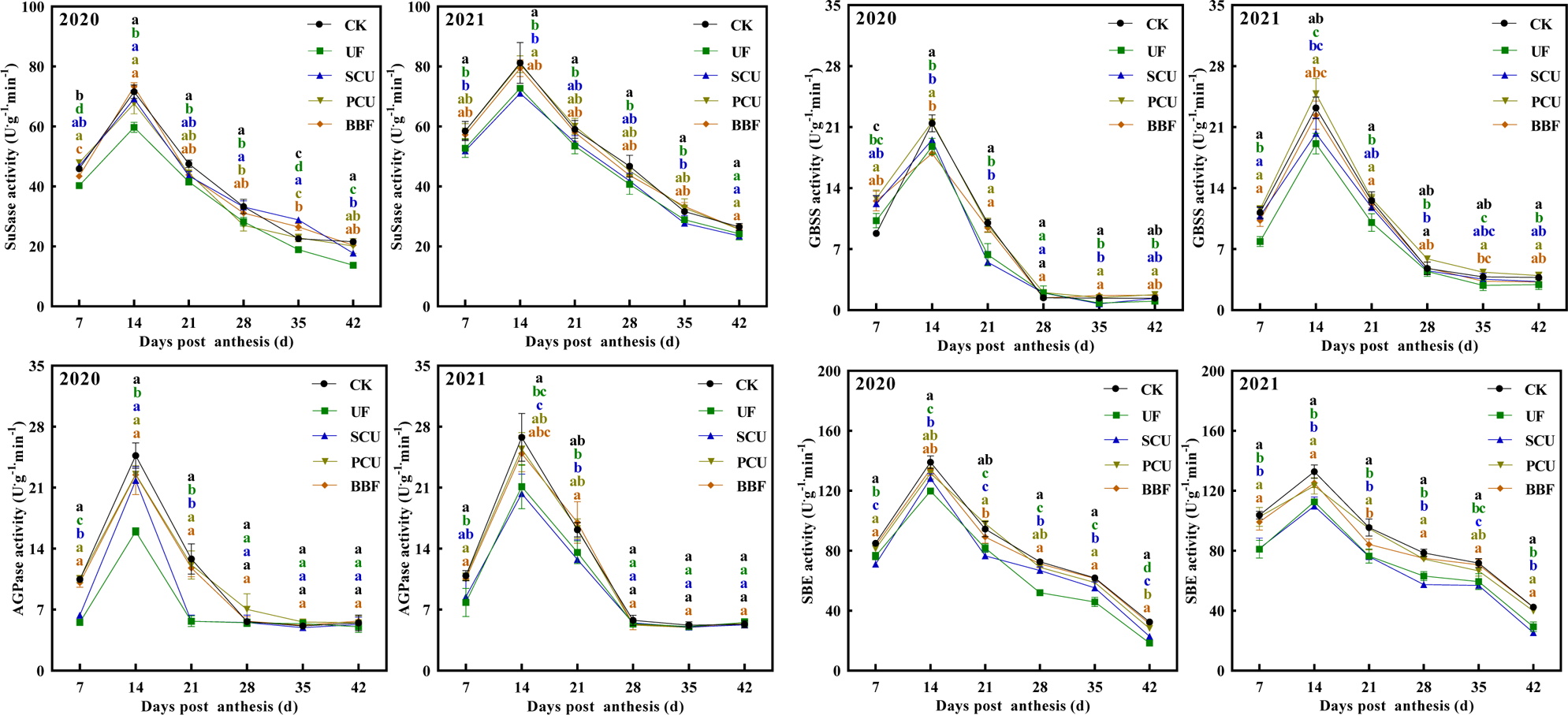
Effects of different types of slow-controlled release fertilizer on the activity of starch synthesis-related enzymes in rice grain during the filling stage in 2020 and 2021. CK, conventional fertilization with four spilt applications of urea; UF, urea formaldehyde; SCU, sulfur-coated urea; PCU, polymer-coated urea; BBF, controlled-release bulk blending fertilizer; Significant differences at each time point are indicated by different letters (*P* < 0.05) as determined by Duncan’s test.

### 3.4 Nitrogen metabolism of rice grain

The formation of amino acids and proteins through nitrogen metabolism process is crucial for the growth of rice grains (Yamakawa and Hakata, 2010). During grain filling, the UF group had significantly lower accumulation of total protein and amino acid in the grain compared to the CK group, whereas the PCU group’s content reached or surpassed the level of CK group in both 2020 and 2021 (Figure. 5). Relative to the CK group, the UF group exhibited significantly lower activities in key enzymes (GS and GOGAT) associated with nitrogen assimilation, whereas the PCU group demonstrated comparable activities in GS and GOGAT to those of the CK group (Figure. 6). During grain filling, the key metabolites and enzyme activities associated with nitrogen metabolism in the grain of both BBF and SCU groups were not significantly higher than those in the PCU group, nor were they lower than those in the UF group (Figures. 5-6).

**Figure 5.**
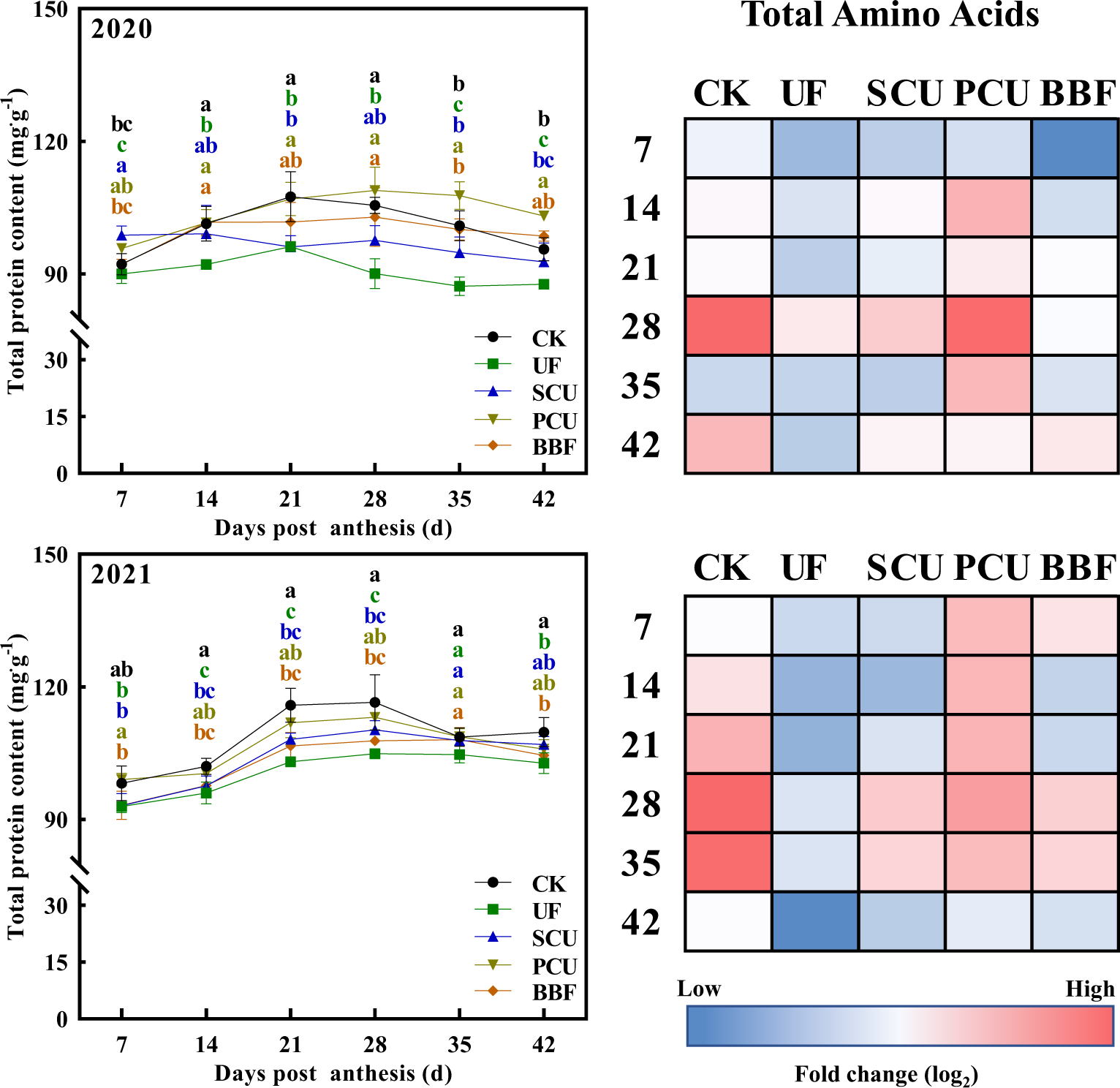
Effects of different types of slow-controlled release fertilizer on the content of total protein and amino acids in rice grain during the filling stage in 2020 and 2021. CK, conventional fertilization with four spilt applications of urea; UF, urea formaldehyde; SCU, sulfur-coated urea; PCU, polymer-coated urea; BBF, controlled-release bulk blending fertilizer; Profiling of total amino acids presented as a heat map calculated by log_2_ fold change (red meaning increased and blue meaning decreased); Significant differences at each time point are indicated by different letters (*P* < 0.05) as determined by Duncan’s test.

**Figure 6.**
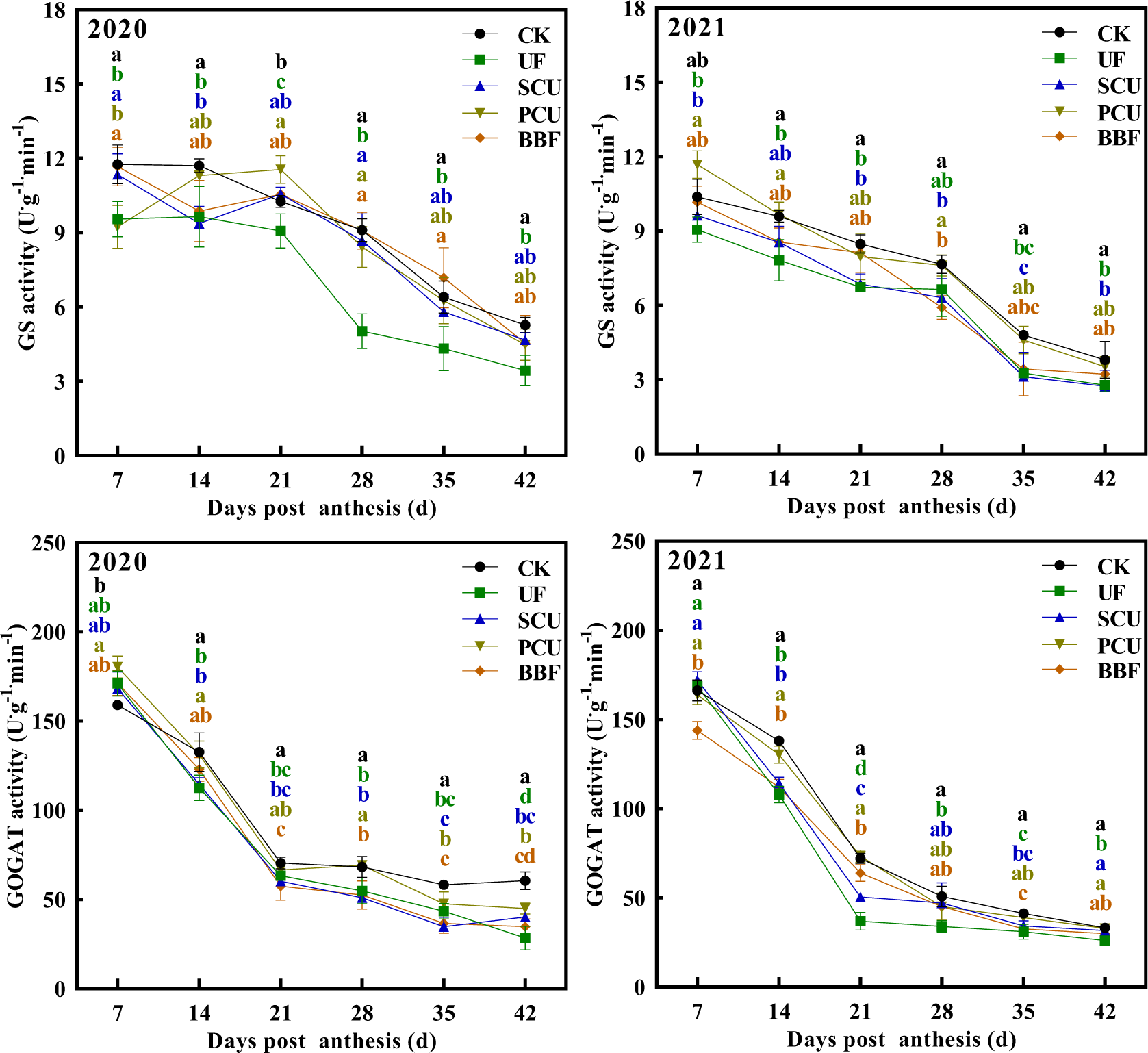
Effects of different types of slow-controlled release fertilizer on changes in nitrogen metabolism-related enzyme activities in rice grain during the filling stage in 2020 and 2021. CK, conventional fertilization with four spilt applications of urea; UF, urea formaldehyde; SCU, sulfur-coated urea; PCU, polymer-coated urea; BBF, controlled-release bulk blending fertilizer; Significant differences at each time point are indicated by different letters (*P* < 0.05) as determined by Duncan’s test.

### 3.5 The crosstalk of carbon and nitrogen metabolism in rice grain

In the dynamic analysis, profiles of sugar-unloading pathway and carbon-nitrogen metabolism were tested during grain filling (Figure. 7). During grain filling, the gene expression of *OsSUT2* and *OsCIN4*, which are related sugar unloading (Ji et al., 2005; Jiang et al., 2021), did not significantly decrease in the groups of UF, PCU and BBF compared to the CK group. This suggested that the application of these fertilizers did not limit the supply of sucrose. The PCU group showed the strongest accumulation of 12 amino acids in rice grain compared to other slow-controlled release fertilizers, and reached the level of the CK group (Figures. 7 and S2). Meanwhile, the UF group showed the lowest accumulation of amino acids among all fertilizer treatments. The accumulation of the BBF group was not higher than that of the PCU group, but not lower than the UF group. The *OsGS2* has a significant impact on both the nitrogen transportation in rice and the metabolism of amino acids in grains (Wang et al., 2022b). During grain filling, the gene expression of *OsGS2* was higher in the groups of PCU and BBF than that of UF and SCU groups. Thus, the increased nitrogen supply in groups of PCU and BBF might enhance nitrogen assimilation and led to an increase in amino acid synthesis of rice grains at the grain filling stage (Table S3, Figures. 1 and S2).

**Figure 7.**
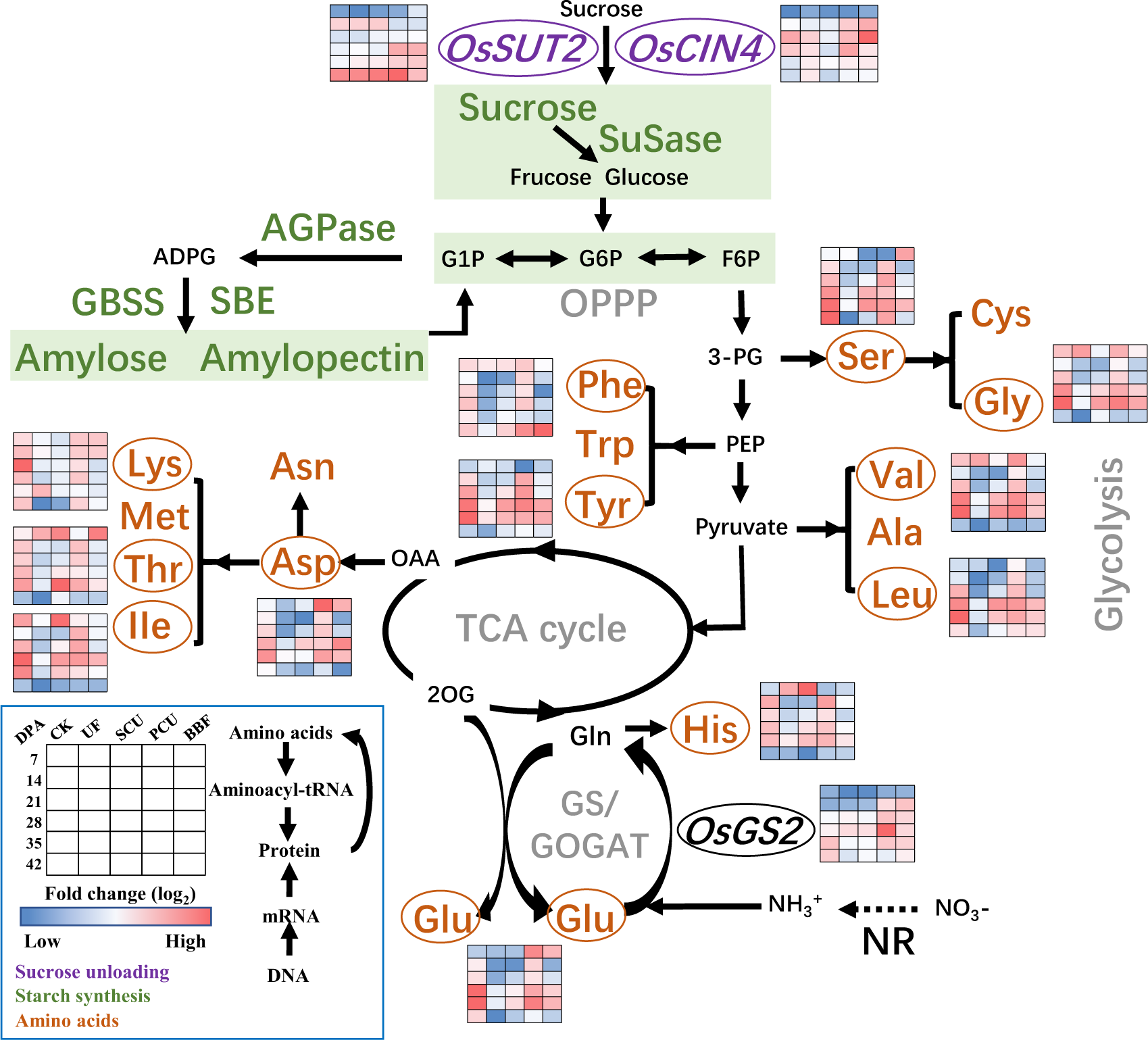
Metabolic profiling corresponding to the accumulation of starch and amino acids in rice grains during the filling stage in 2021. CK, conventional fertilization with four spilt applications of urea; UF, urea formaldehyde; SCU, sulfur-coated urea; PCU, polymer-coated urea; BBF, controlled-release bulk blending fertilizer; DPA, days post anthesis; Values of the heat map calculated by log_2_ fold change (red meaning increased and blue meaning decreased).

## 4. Discussion

### 4.1 Rice yield and quality are improved by the application of optimal slow-controlled release fertilizer

Compared to the conventional fertilizer, the UF and SCU groups exhibited a significantly lower number of spikelets per panicle and area, which notably inhibited rice yield (Table 1). Applying an optimal nitrogen source close to the grain-filling stage can effectively promote yield formation (Nasielski and Deen, 2019). Compared to the UF and SCU fertilizers, the PCU and BBF fertilizers consistently released nitrogen throughout the growth stage, providing more nitrogen for panicle differentiation and grain filling (Figure. 1). Intriguingly, the rice yield of PCU and BBF groups did not show significant differences compared to CK group, and the application of slow-controlled release fertilizers obviously improved grain weight (Table 1, Figure 2). In 2020, the BBF group achieved the highest yield of 10.8 t hm^−2^ among all fertilizers (Table 1). In 2021, the BBF yield increased to 11.0 t hm^−2^, significantly surpassing the UF and SCU groups’ yields (Table 1).

Variations in nitrogen supply significantly impact the distribution and buildup of starch and protein during grain filling, ultimately affecting grain quality (Wang et al., 2021b; Xu et al., 2021). Compared to conventional fertilizer, the application of UF and SCU fertilizers led to a significant reduction in nutritional value, while the PCU group exhibited lower cooking and eating quality (Tables 2-3). The low nutritional value observed in the UF and SCU groups might be attributed to insufficient nitrogen supply during the grain filling stage (Figure. 1), based on the report that inadequate nitrogen reduced synthesis of protein and amino acids (Wang et al., 2021b). In addition, the cooking and eating quality of rice can be adversely reduced by high nitrogen application (Shi et al., 2023), the low quality of the PCU group might be attributed to the release of excess nitrogen during the grain filling stage (Table 2, Figure. 1). Conversely, the application of BBF fertilizer led to good grain quality in terms of physical appearance, cooking and eating qualities, and nutritional value (Tables 2-3). This was achieved due to its optimal nitrogen release during grain filling (Figure. 1), with each quality-related index reaching those of CK group and surpassing other fertilizer treatments (Tables 2-3). Hence, to ensure a high rice yield and quality, it is crucial to employ an optimal slow-controlled release fertilizer, such as BBF.

### 4.2 The carbon and nitrogen balance in rice grains is influenced by the type of slow-controlled release fertilizer

The accumulation of starch, protein, and amino acids determines grain quality, and this process requires significant amounts of carbon and nitrogen metabolites (Yamakawa and Hakata, 2010; Jiang et al., 2021; Liu et al., 2021). The application of slow-controlled release fertilization did not hinder sucrose accumulation but led to a decrease in starch and amylopectin content in the PCU group compared to the CK group (Table 2, Figure. 3). Meanwhile, the PCU group showed no reduction in enzyme activity related to carbon metabolism (Figure. 4). In contrast, the UF group decreased the activities of SuSase, AGPase, and SBE, but did not show lower starch and amylopectin content compared to the PCU group (Table 2, Figures. 3-4). Carbon metabolism is crucial in nitrogen metabolism as it provides the carbon skeletons necessary for amino acid synthesis, including pyruvate, oxaloacetate, and alpha-ketoglutarate (Xu et al., 2012). Thus, the complex results observed in this study might be linked to the interplay between carbon and nitrogen metabolism (Baunsgaard et al., 2005; Seung et al., 2013; MacNeill et al., 2017).

The PCU and BBF fertilizers had a higher nitrogen-release ability during grain filling compared to UF and SCU fertilizers (Ye et al., 2013; Wu et al., 2021). Intriguingly, the application of UF resulted in a significant reduction in accumulation of protein and amino acids compared to conventional fertilizer, whereas PCU fertilizers achieved levels similar to those of the CK group during grain filling (Figure. 5). The GS/GOGAT cycle plays a key role in nitrogen assimilation (Gao et al., 2019; Hou et al., 2019). During grain filling, the PCU group showed increased activity in GS and GOGAT enzymes, reaching levels comparable to those of the CK group, whereas the UF group showed significantly lower activity in these enzymes (Figures. 5-6). These results indicated that increased nitrogen supply mainly leads to increased synthesis of protein and amino acids in rice grains rather than increased accumulation of storage starch. However, the BBF group showed a better partition in starch and protein content, which was neither the highest nor the lowest compared to others (Figures. 3-6). This phenomenon may be related to the appropriate nitrogen release ability of BBF fertilizer during grain filling (Figure. 1). Despite the complex relationship between protein production and carbon-nitrogen metabolism, the dynamic changes in protein synthesis are significantly related to the application of slow-controlled release fertilizers (Table S3, Figure. S2).

### 4.3 Key steps for promoting carbon-nitrogen crosstalk in developing grain under the application of slow-controlled release fertilizer

Maintaining a balance between carbon and nitrogen metabolism is crucial for grain filling of rice, as nitrogen, functioning as a signaling molecule, influences plant metabolism and physiology through changes in gene expression (Xin et al., 2019). During the grain-filling stage, rice grain undergoes dynamic metabolic adjustments to meet its nutritional demands (Jiang et al., 2022a). An analysis of the primary changes in amino acid metabolism revealed clear differences in the carbon and nitrogen metabolism among various slow-controlled release fertilizers (Figure. 7).

The expression of genes (*OsSUT2* and *OsCIN4*) related to sucrose unloading (Sekhar et al., 2021), as well as sucrose content, in the UF, PCU, and BBF groups were not lower than that of CK group (Figure. 3 and 7), suggesting that the availability of carbohydrates was not restricted by these application of slow-controlled release fertilizers. N-nutrient/metabolites and carbon metabolism are intricately interconnected (Yamakawa and Hakata, 2010). Carbon metabolism plays a critical role in incorporating nitrogen into cell metabolism, primarily through the oxidative pentose phosphate pathway (OPPP) and glycolysis/ the tricarboxylic acid (TCA) cycle (Bussell et al., 2013; Yang et al., 2016; Wang et al., 2021a; Jiang et al., 2022b). When comparing various types of slow-controlled release fertilizers, it was found that the application of PCU and BBF fertilizers led to a higher capacity for sucrose unloading and nitrogen assimilation (Figures. 7 and S2). In the metabolic analysis, the increased carbon flux was primarily directed towards the glycolysis and TCA cycle for amino acid synthesis in the PCU and BBF groups (Figure. 6 and S2). As the increase of nitrogen supply in PCU and BBF (Figure. 1), the GS/GOGAT cycle was significantly promoted in rice grain at the filling stage (Table S2, Figures. 7 and S2). The PCU group demonstrated significantly higher enzyme activity in carbon and nitrogen metabolism and a lower starch content in rice grain, compared to the UF and SCU groups (Figures. 3-6). A high supply of nitrogen can increase the sugar unloading and metabolic utilization (Zhang et al., 2021). Consequently, the PCU group’s high capacity for nitrogen assimilation lead to efficient utilization of carbohydrates for amino acid synthesis during grain filling. Interestingly, the BBF group showed a relatively balanced performance in key enzymes of carbon and nitrogen metabolism, as well as in the levels of starch, protein, and amino acids (Figures. 3-6). These results suggest that a suitable supply of nitrogen promotes both grain filling and maintains an optimal carbon-nitrogen state in the rice grain. Therefore, the application of slow-controlled release fertilizer like BBF is a practical solution for manipulating the balance between carbon and nitrogen in rice.

Overall, these observations suggest that an increase in nitrogen supply not only enhances nitrogen metabolism in grain, but also improves carbon flow (Table S2, Figures. 7 and S2). Our study proposes a model in which nitrogen supply regulates the networks of carbon-nitrogen metabolism, and the regulation of starch and protein synthesis under high nitrogen conditions is linked to nitrogen assimilation and the glycolysis/TCA cycle (Table S3, Figure. 7 and S2).

### 4.4 Conclusions

The application of BBF fertilizer not only increased rice yield but also enhanced grain quality compared to other slow-release fertilizers, such as UF, SCU, and PCU. Moreover, it has the potential to reach or even exceed the levels achieved with conventional fertilization. This can be primarily attributed to BBF fertilizer’s superior regulation of the carbon-nitrogen balance in rice grains compared to other slow-controlled release fertilizers. The application of BBF fertilizer appropriately increases nitrogen assimilation (amino acid synthesis and nitrogen transport) in the grain by modulating carbon flow in the carbon metabolism of grain (e.g., glycolytic metabolism and the TCA cycle). Our research provides new insight into the relationship among the nitrogen supply, metabolism pathway of carbon and nitrogen, and grain growth to finely promote grain quality under the application of slow-controlled release fertilizer in rice.

## Abbreviations

CK: Conventional fertilization
UF: urea formaldehyde fertilizer
SCU: sulfur-coated urea fertilizer
PCU: polymer-coated urea fertilizer
BBF: controlled-release bulk blending fertilizer
TCA: the tricarboxylic acid
NUE: nitrogen use efficiency
GS: Glutamine synthetase
GOGAT: glutamate synthase
OPPP: oxidative pentose phosphate pathway
SuSase: Sucrose Synthase
AGPase: ADP-glucose pyrophosphorylase
GBSS: granule-bound starch synthase
SBE: starch branching enzyme
Glu: glutamine
Gly: glycine
His: histidine
Ile: isoleucine
Leu: leucine
Lys: lysine
Phe: phenylalanine
Ser: serine
Thr: threonine
Tyr: tyrosine
Val: valine
DPA: Days post-anthesis
RVA: rapid viscosity analyzer
DW: Dry weight.

## Declarations

### Ethics approval and consent to participate

Not applicable

### Availability of Supporting Data

The datasets generated and/or analysed during the current study are available from the corresponding author on reasonable request.

### Author Contributions

ZJ and QC: Conceptualization, Software, Formal analysis, Investigation, Writing - Original Draft and Visualization; DL, WT, SG, and JL: Validation and Investigation. CL and MZ: Data Curation; YD, and WL: Resources and Supervision; GL and SS: Conceptualization and Revising - Original Draft; LX: Draft revision, Project administration and Funding acquisition. All authors read and approved the final manuscript.

## Acknowledgments

The authors thank Maria-Dolores Pérez-Garcia, and Laurent Ogé for kindly providing guidance and Gaoya Sun for providing technical assistance.

## Supporting Information

**Table S1.**
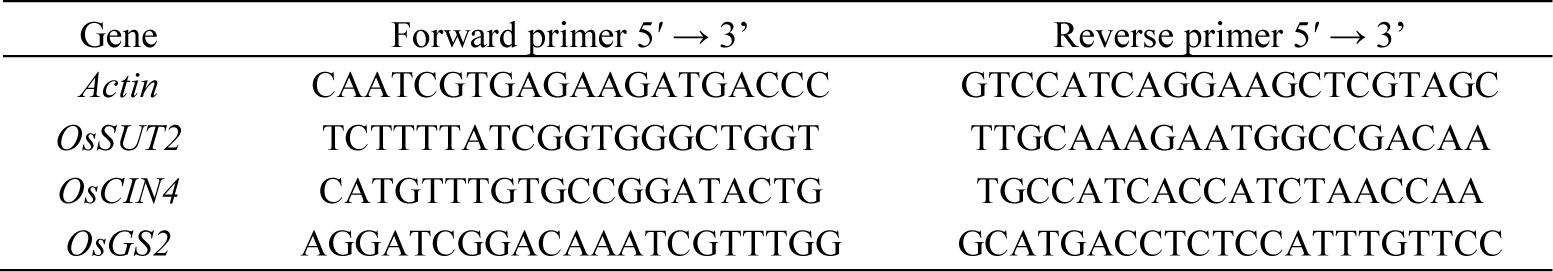
Sequences of primers for Actin and genes for qRT-PCR.

**Table S2.**
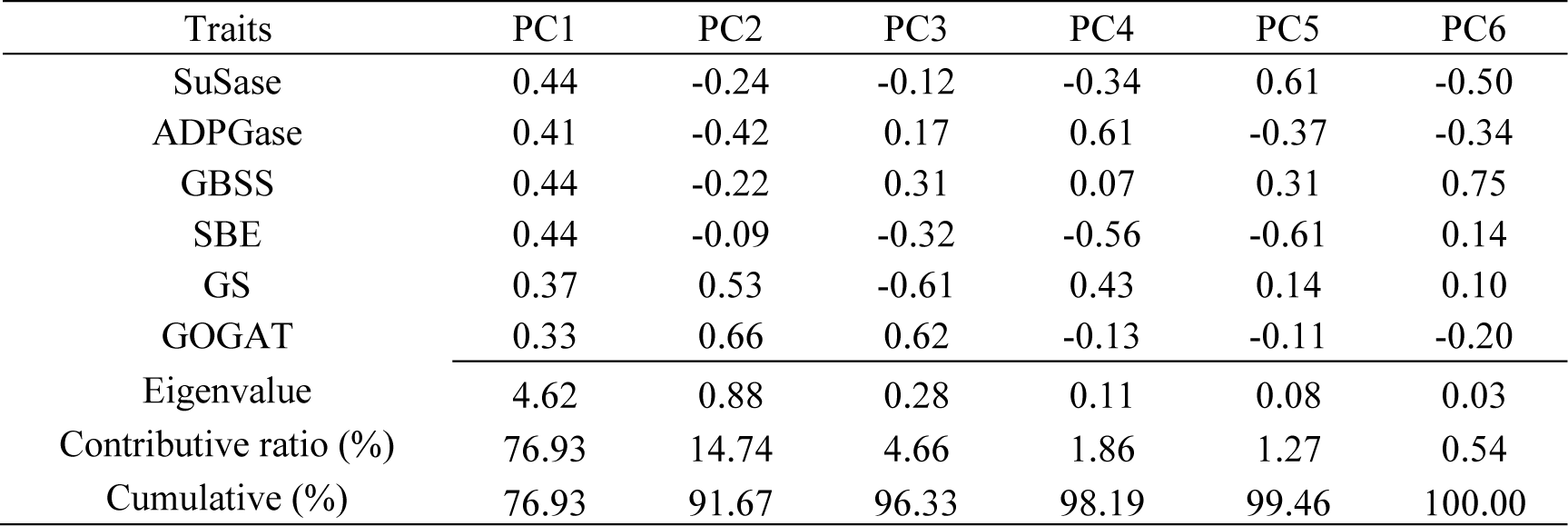
Principal components analysis about the effect of slow-controlled release fertilizer on key enzymes activities related carbon-nitrogen metabolism in rice grain at filling stage in 2020 and 2021.

| Traits | PC1 | PC2 | PC3 | PC4 | PC5 | PC6 |
| --- | --- | --- | --- | --- | --- | --- |
| SuSase | 0.44 | -0.24 | -0.12 | -0.34 | 0.61 | -0.50 |
| ADPGase | 0.41 | -0.42 | 0.17 | 0.61 | -0.37 | -0.34 |
| GBSS | 0.44 | -0.22 | 0.31 | 0.07 | 0.31 | 0.75 |
| SBE | 0.44 | -0.09 | -0.32 | -0.56 | -0.61 | 0.14 |
| GS | 0.37 | 0.53 | -0.61 | 0.43 | 0.14 | 0.10 |
| GOGAT | 0.33 | 0.66 | 0.62 | -0.13 | -0.11 | -0.20 |
| Eigenvalue | 4.62 | 0.88 | 0.28 | 0.11 | 0.08 | 0.03 |
| Contributive ratio (%) | 76.93 | 14.74 | 4.66 | 1.86 | 1.27 | 0.54 |
| Cumulative (%) | 76.93 | 91.67 | 96.33 | 98.19 | 99.46 | 100.00 |

**Table S3.** Principal components analysis of comprehensive index and loading matrix of each component for amino acids accumulation under the application of slow-controlled release fertilizers in 2020 and 2021.

| Traits | PC1 | PC2 | PC3 | PC4 | PC5 | PC6 | PC7 | PC8 | PC9 | PC10 | PC11 | PC12 |
| --- | --- | --- | --- | --- | --- | --- | --- | --- | --- | --- | --- | --- |
| Asp | 0.30 | 0.13 | 0.14 | -0.49 | 0.22 | -0.16 | 0.35 | 0.28 | -0.16 | -0.57 | -0.08 | -0.05 |
| Thr | 0.28 | -0.39 | 0.08 | 0.08 | 0.00 | 0.62 | -0.09 | 0.17 | -0.50 | -0.04 | 0.30 | 0.06 |
| Ser | 0.25 | 0.31 | -0.47 | 0.19 | 0.21 | 0.48 | 0.17 | 0.08 | 0.50 | -0.13 | 0.05 | 0.06 |
| Glu | 0.34 | 0.26 | -0.02 | -0.24 | 0.17 | 0.04 | 0.28 | -0.07 | -0.25 | 0.73 | -0.21 | 0.01 |
| Gly | 0.35 | -0.08 | 0.09 | -0.21 | -0.09 | 0.17 | -0.32 | -0.49 | 0.16 | -0.11 | -0.22 | -0.59 |
| Val | 0.36 | -0.14 | 0.12 | -0.20 | -0.10 | -0.04 | -0.27 | -0.20 | 0.23 | -0.04 | -0.17 | 0.77 |
| Ile | 0.32 | -0.32 | 0.09 | 0.01 | 0.06 | -0.26 | -0.19 | 0.61 | 0.40 | 0.28 | 0.13 | -0.22 |
| Leu | 0.30 | 0.39 | -0.15 | -0.04 | -0.28 | -0.29 | -0.15 | -0.15 | -0.12 | -0.01 | 0.71 | -0.01 |
| Tyr | 0.27 | -0.02 | -0.55 | 0.31 | -0.24 | -0.26 | -0.16 | 0.17 | -0.36 | -0.13 | -0.46 | -0.02 |
| Phe | 0.13 | 0.45 | 0.56 | 0.31 | -0.48 | 0.19 | 0.04 | 0.24 | 0.04 | -0.04 | -0.21 | -0.02 |
| Lys | 0.20 | 0.14 | 0.29 | 0.52 | 0.67 | -0.21 | -0.20 | -0.17 | -0.13 | -0.09 | 0.02 | 0.02 |
| His | 0.26 | -0.40 | 0.08 | 0.33 | -0.21 | -0.18 | 0.68 | -0.31 | 0.13 | -0.03 | 0.08 | -0.02 |
| Eigenvalue | 6.57 | 1.47 | 0.99 | 0.89 | 0.74 | 0.47 | 0.29 | 0.19 | 0.19 | 0.08 | 0.07 | 0.04 |
| Contributive ratio (%) | 54.75 | 12.25 | 8.29 | 7.44 | 6.19 | 3.93 | 2.45 | 1.59 | 1.55 | 0.70 | 0.55 | 0.32 |
| Cumulative (%) | 54.75 | 67.00 | 75.29 | 82.73 | 88.92 | 92.84 | 95.29 | 96.89 | 98.44 | 99.13 | 99.68 | 100.00 |

**Figure S1.**
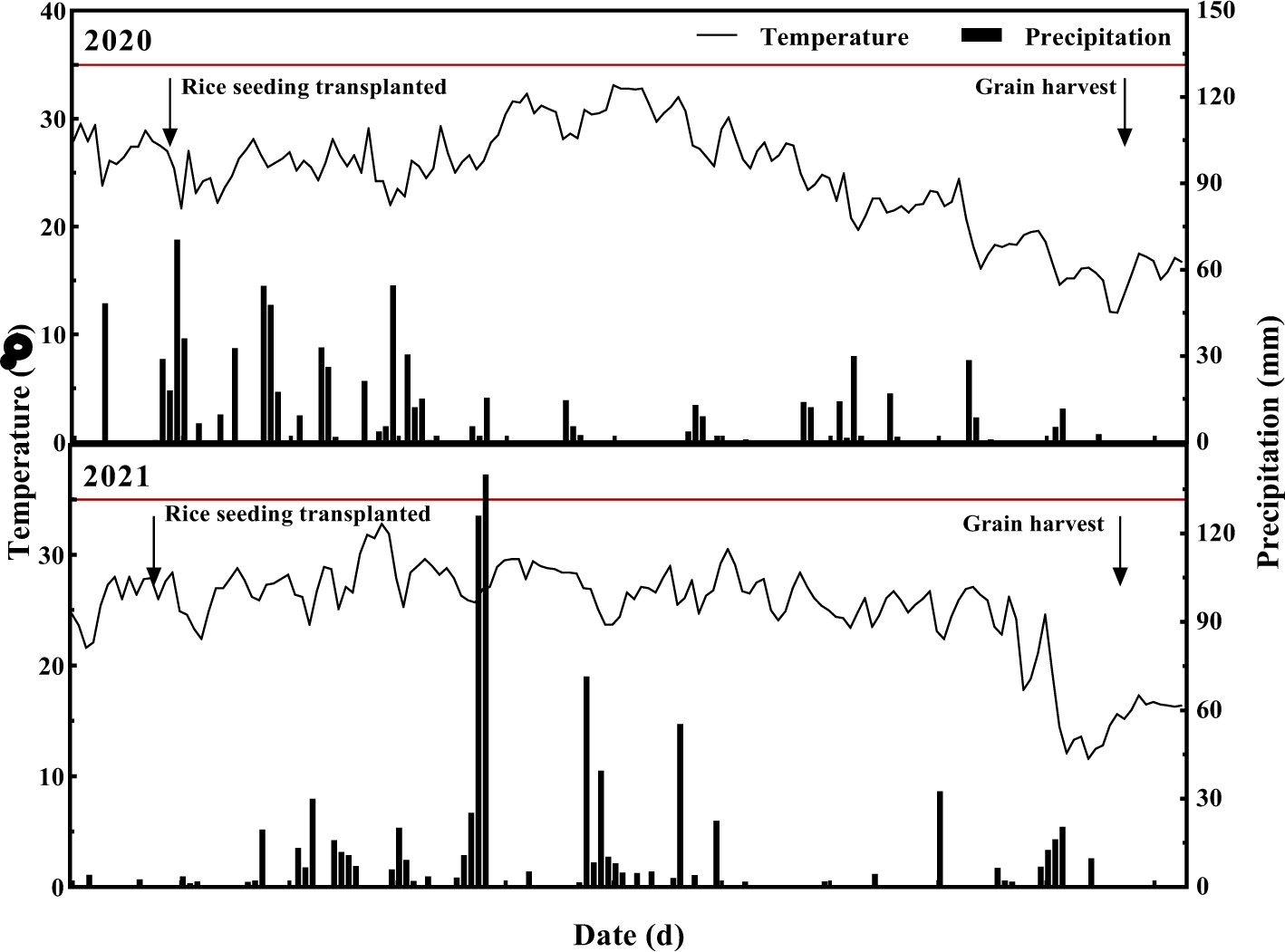
Daily temperature (line) and precipitation (bars) during the rice growth seasons in 2020 and 2021.

**Figure S2.**
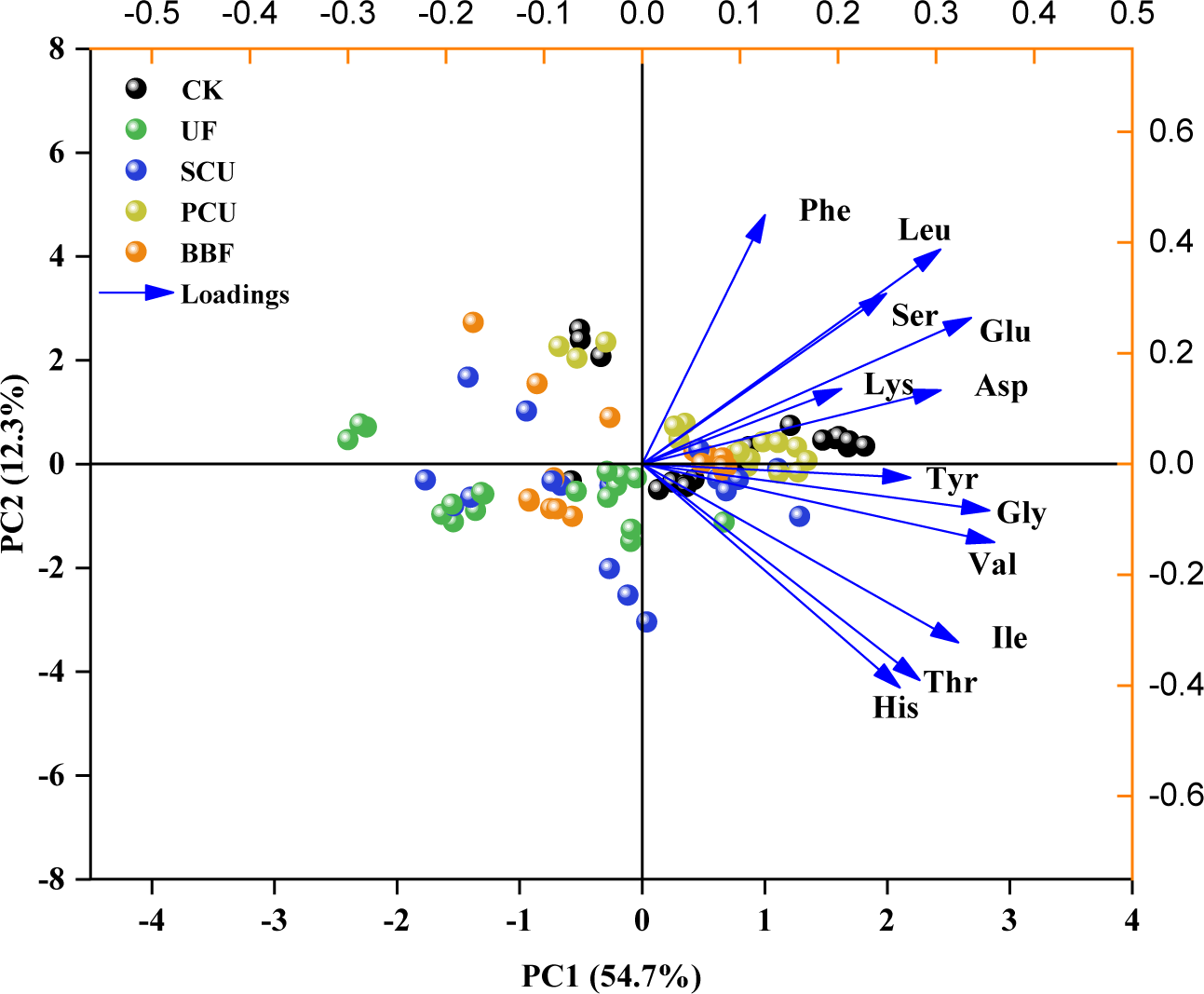
Principal component analysis about the effect of slow-controlled release fertilizer application on the accumulation of amino acids in rice grains at the filling stage in 2020 and 2021. CK, conventional fertilization with four spilt applications of urea; UF, urea formaldehyde; SCU, sulfur-coated urea; PCU, polymer-coated urea; BBF, controlled-release bulk blending fertilizer.

